# Alpha satellite RNA marks the perinucleolar compartment and represses ribosomal RNA expression in naïve human embryonic stem cells

**DOI:** 10.1101/2025.07.11.664446

**Authors:** Kirti Mittal, Lamisa Ataei, Miguel Ramalho-Santos

**Affiliations:** Lunenfeld-Tanenbaum Research Institute, Mount Sinai Hospital, Toronto, Ontario M5G 1X5, Canada; Department of Molecular Genetics, University of Toronto, Toronto, Ontario M5S 1A8, Canada

## Abstract

While most newly synthesized RNA is exported to the cytoplasm, a portion of non-coding RNA is retained in the nucleus and remains highly associated with chromatin. The strong binding of this RNA fraction to insoluble chromatin impairs its recovery in standard transcriptomic studies. Therefore, the landscape and potential functions of chromatin-associated RNAs are poorly understood. Recent studies indicate that chromatin-associated transcripts can have regulatory roles, particularly during mammalian development. Here we compare the dynamics of cytoplasmic versus chromatin-bound transcriptomes of naïve and primed human embryonic stem cells (hESCs), as well as fibroblasts. We found a remarkable enrichment for RNA transcribed from alpha satellite repeat (ALR) in the chromatin fraction of naïve hESCs, compared to primed hESCs. The co-localization and interaction of ALR RNA with Polypyrimidine Tract Binding Protein 1 (PTBP1) protein indicate that ALR RNA foci mark the perinucleolar compartment (PNC), a nuclear sub-compartment previously thought to be exclusive to cancer cells. Knockdown of ALR RNA leads to dispersion of PTBP1 foci, up-regulation of ribosomal RNA and global hypertranscription in naïve hESCs. In contrast, loss of PTBP1 does not disturb ALR RNA localization, indicating that ALR is upstream in the hierarchy of organization of the PNC in hESCs. These results reveal a role for ALR RNA in nuclear compartmentalization and tuning rRNA synthesis in naïve hESCs. Moreover, this study opens new avenues to dissect the function of ALR RNA and the PNC in cancer contexts.

## INTRODUCTION

The transcriptome represents the portion of the genome that is actively expressed in any given cell. Thus, transcriptomic methods such as RNA-sequencing (RNA-seq) provide a powerful lens to study cellular state dynamics during development, adult physiology and disease. Standard RNA-seq methods focus on cytoplasmic RNA, which primarily represents cytosolic transcripts. However, it is becoming increasingly clear that there exists a “hidden” transcriptome tightly bound to chromatin. Because of its poor solubility, this fraction of chromatin-associated RNAs (caRNAs) is often overlooked in typical RNA-seq methods^1–4^. Recent methods involving first fractionating cells to isolate insoluble chromatin and subsequently extracting RNA for deep sequencing have begun to shed light on the nature and possible functions of caRNAs^5–8^.

CaRNAs are typically non-coding and enriched in repeat sequences. Functional studies to date support the notion that caRNAs play roles in higher-order chromosomal architecture^1,5,9–17^. For example, the well-studied lncRNA *XIST* coats one of the X chromosomes and promotes transcriptional silencing during X chromosome inactivation^18–21^. Nascent, repeat-rich RNA has been shown to associate strongly with chromatin and contribute to maintenance of decondensed chromosome territories^22^. Long interspersed nuclear element 1 (LINE1) is one of the most abundant classes of repeats in both mouse and human genomes. LINE1 RNA binds to chromatin, maintains nucleolar architecture and prevents developmental reversion in mouse and human embryonic stem cells^9,23–25^. Recently, the chromatin-bound transcriptomes of primed hESCs and other cell lines were characterized, revealing a global interdependence between caRNAs and 3D genome organization^2,3,26^. These and other studies highlight the potential for genomic regulatory information that resides in the chromatin-bound transcriptome, particularly during early development.

In this study, we optimized a methodology to successfully isolate caRNAs from two types of pluripotent cells, naïve and primed hESCs, using fetal lung fibroblasts as somatic cell controls. Standard cultures of hESCs maintain them in a primed state, which corresponds to a post-implantation stage of pluripotent cells in the human embryo. The naïve state in vitro captures an earlier, pre-implantation stage of pluripotency in vivo^27–29^. We report that alpha satellite repeat (ALR) RNA is highly enriched in naïve but not primed hESCs. This result aligns well with an upregulation of ALR RNA at the morula/blastocyst stage of human development, prior to implantation^30^. ALR RNA in naïve hESCs is present in highly defined foci in the vicinity of the nucleolus and co-localized with Polypyrimidine Tract Binding Protein 1 (PTBP1). PTBP1 is a marker of the perinucleolar compartment (PNC), previously described in cancer cells^31,32^. We found that ALR RNA is required for maintenance of the PNC in naïve hESCs. Our functional studies further indicate that ALR RNA contributes to maintenance of low levels of ribosomal RNA expression and global transcriptional output of naïve hESCs.

## RESULTS

### The chromatin-bound transcriptome of hESCs is enriched for transcripts derived from genomic repeats

We aimed to define the chromatin-bound transcriptome of different states of pluripotency in hESCs. We developed a methodology based on cell fractionation protocols previously described in other cell types^7,8^ and optimized it for the enrichment of the chromatin fraction of hESCs (see Methods and Supplemental Fig. 1A). Cell types used were naïve hESCs (cultured in RSeT medium^33^), primed hESCs (cultured in mTeSR medium^34^) and fetal lung fibroblasts (IMR-90 cell line) (Fig. 1A). While there are various formulations for culture of naïve and primed hESCs, the media used afforded the reproducibility of standardized commercial formulations, given the need to grow large numbers of cells for fractionation. Moreover, we have validated the RSeT base medium in other recent studies of the naïve hESC state^24,25^, and confirmed key findings of this study in alternative naïve medium formulations (see below, Supplemental Fig.4D). Validation of the fractionation strategy was carried out using RT-qPCR for markers of mature mRNAs (cytoplasmic) vs nascent pre-mRNA and pre-rRNA (chromatin-bound) (Supplemental Fig. 1A).

**FIGURE 1.**
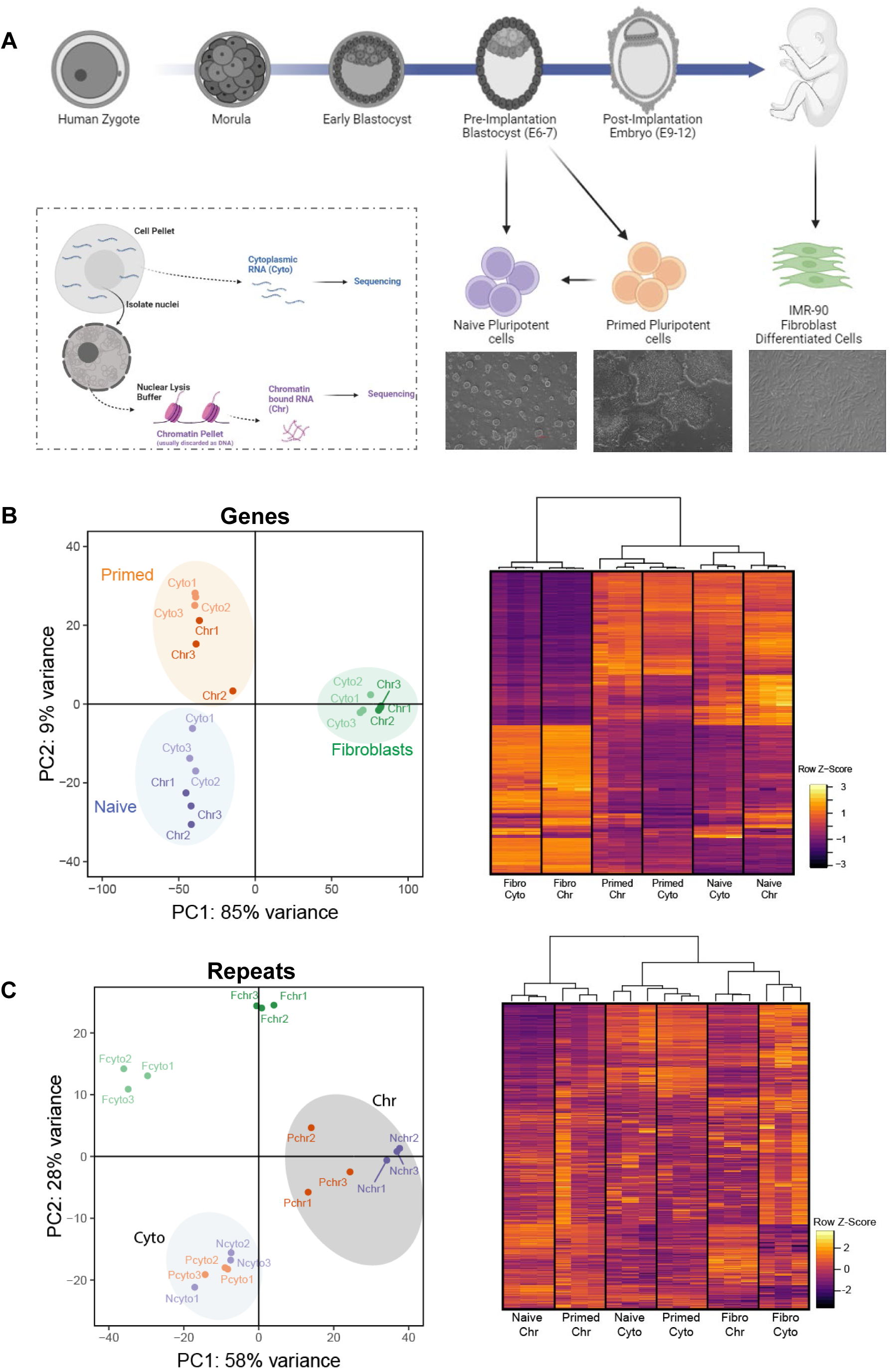
Fractionation and identification of cytoplasmic and chromatin-bound RNA reveals a distinct profile of repeat-derived transcripts in hESCs. A) Schematic of early human embryonic development with the cell types used for cell fractionation and RNA-Seq. B) PCA plot for all expressed genes across all samples, showing across PC1 that fibroblasts acquire a more divergent expression profile as compared to the Naïve and Primed hESCs; Hierarchical clustering of top 2000 most variable genes in all cell types shows clustering by cell type (n = 3 biological replicates per group). C) PCA plot for all expressed repeats across all samples, showing across PC1 that chromatin fractions acquire a more divergent expression profile as compared to cytosolic fractions; Hierarchical clustering of all repeats expressed shows clustering by fraction type (n = 3 biological replicates per group).

We then performed RNA-sequencing from three biological replicates of the cytosolic and chromatin-associated fractions from all these cell types (Fig. 1A). We first confirmed that markers of the naïve vs primed hESC state^35,36^ are indeed enriched in their respective samples (Supplemental Fig. 1B,C). Principal component analysis (PCA) of unique, protein-coding genes revealed a strong similarity between the cytoplasmic and the chromatin-bound fractions of each cell type, and a clear separation between each of the three cell types. Thus, the difference in the cell types is the dominant factor of variation of the protein-coding transcriptome, regardless of fraction (Fig. 1B). Surprisingly, PCA specifically of transcripts emerging from repeat sequences showed that the chromatin fractions across cell types diverge from their respective cytosolic fractions (Fig. 1C). Hierarchical clustering yields similar results, with a notable distinct clustering of naïve and primed chromatin-bound repeat transcriptome (Fig. 1B, C).

We focused our attention on the chromatin-bound transcriptome. We validated that known chromatin-bound lncRNAs (XIST, KCNQ1OT1, NEAT1, MALAT1) and cytosolic mature mRNAs (ACTB, GAPDH) are enriched in the expected fraction in all cell types (Fig. 2A). As expected, RNA from introns, which are present in chromatin-associated nascent transcripts but are spliced out before nuclear export, are also present in the chromatin fraction (Fig. 2B). Interestingly, the chromatin fraction is highly enriched for transcripts of repeat sequences, across all cell types (Fig. 2C). Taken together, these findings point to distinct dynamics of the protein-coding transcriptome versus the repeat transcriptome across different cell types and suggest that repeat-derived caRNAs may have novel roles in hESCs.

**FIGURE 2.**
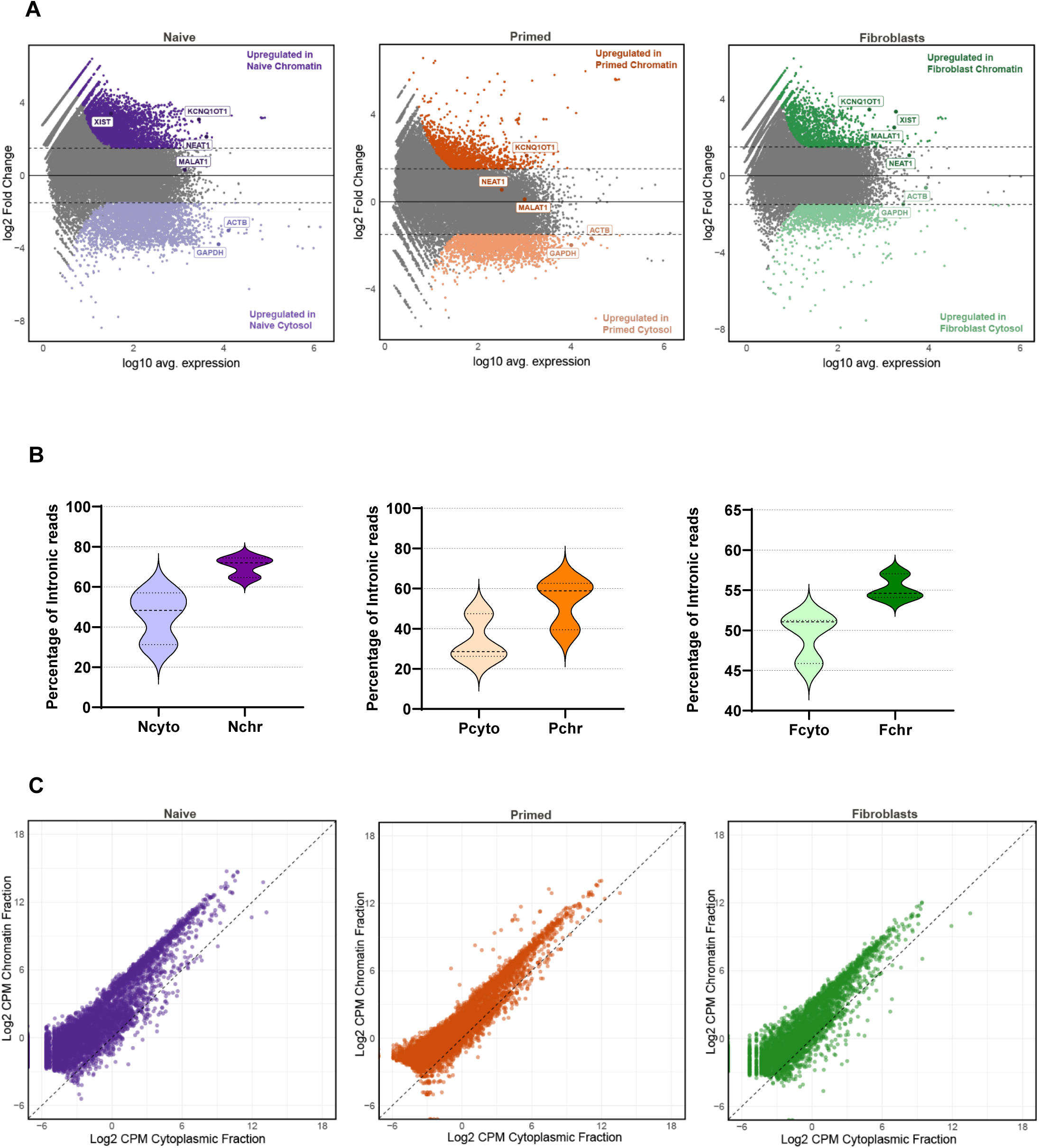
The chromatin-associated transcriptome is highly enriched for repeat-derived RNA. A) Comparison of chromatin versus cytoplasmic transcriptomes across cell types. Indicated are known chromatin associated lncRNAs (XIST, KCNQ1OT1, MALAT and NEAT1) and cytoplasmic mRNAs (GAPDH, ACTB). B) Intronic reads are higher in the chromatin fractions from all cell types compared to the cytosolic fraction. C) Scatter plots showing that repeat-derived RNAs are highly enriched in the chromatin fraction, for all the 3 cell types. CPM, counts per million.

### ALR RNA is a specific, highly expressed chromatin-associated transcript in naïve hESCs

We next sought to identify repeat-derived transcripts that distinguish the naïve and primed states of human pluripotency. Differential expression analyses revealed a remarkable up-regulation of alpha satellite ALR in naïve hESCs, relative to primed cells (Fig. 3A, Supplemental Fig. 2A,2B, 2C). Moreover, ALR RNA is strongly enriched at the chromatin of both naïve and primed hESCs (Fig. 3B, Supplemental Fig. 2D).

**FIGURE 3.**
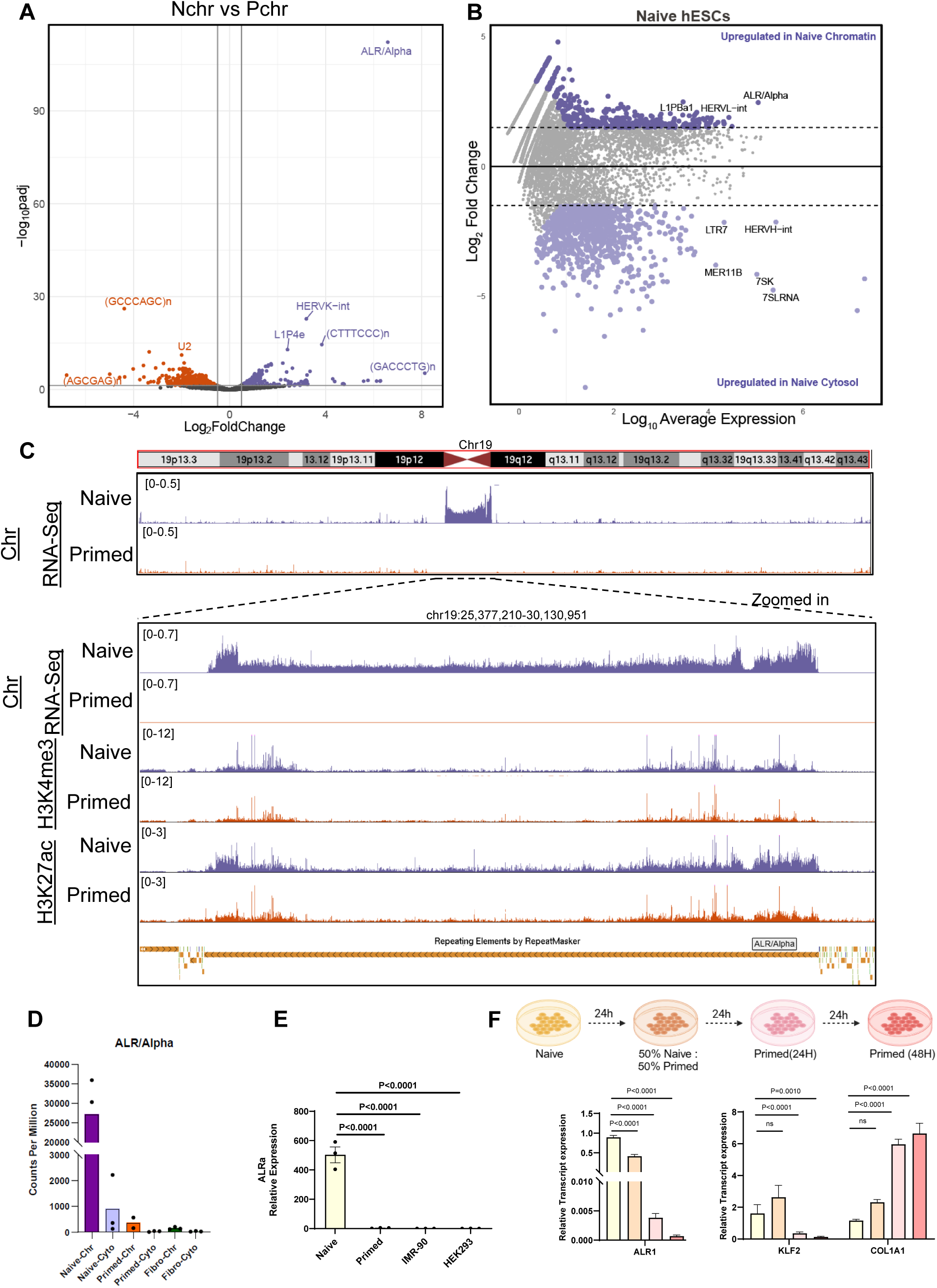
ALR RNA is a specific, highly expressed chromatin-associated transcript in naïve hESCs. A) Volcano plot comparing Naïve versus Primed chromatin fractions, with strong enrichment of ALR repeats in naïve chromatin. B) MA plot comparing Naïve chromatin versus Naïve cytosolic transcriptomes, showing abundant expression and enrichment of ALR in the chromatin fraction. C) Genome browser view of chromosome 19, focusing on the ALR-rich centromere. Chromatin-bound RNAseq and ChIP-seq tracks for histone marks in naïve versus primed hESCs are shown (see the Materials and Methods). Tracks from each sequencing experiment were normalized to the same scale between cell types. D) Expression level (CPM) of ALR, showing very high expression in the chromatin fraction of Naïve hESCs. The expression of ALR at chromatin of each cell type is higher than their cytosolic counterparts. E) Detection of ALR expression in naïve hESC, primed hESC, embryonic fibroblasts (IMR-90) and HEK293T cells by RT-qPCR. Data are average fold change relative to Naïve hESCs ± SEM, n = 3 biological replicates. The statistical significance of the difference between naïve hESCs and other cell types was determined by one-way ANOVA and Dunnett’s multiple comparisons test. F) Conversion of naïve hESCs towards the primed state shows a sharp decrease in relative expression of ALR and naïve markers and increase in primed markers. The statistical significance of the difference between naïve hESCs and other cell types was determined by one-way ANOVA and Dunnett’s multiple comparisons test.

ALR is a satellite repeat enriched at the centromeric and pericentromeric regions of all chromosomes. Because of the highly repetitive nature of these regions, they have been missed in previous assemblies of human genome, complicating data analysis. The recent T2T human genome assembly (build -CHM13v1.1) has annotated centromeric and pericentromeric regions to a much better extent^37^. Upon alignment of the RNA-seq data to the T2T genome assembly, we observed a robust signal of ALR RNA at the centromeric and pericentromeric regions of naïve hESCs, with minimal to no detection in primed hESCs or fibroblasts (Fig. 3C, and data not shown). We also mapped onto the T2T genome ChIP-Seq data sets for histone marks in naïve and primed hESCs, generated in partnership with the Canadian Epigenetics, Environment, and Health Research Consortium (CEEHRC). This analysis revealed a clear enrichment of active histone marks H3K4me3 and H3K27ac at ALR repeats in naïve hESCs (Fig. 3C). In contrast, the repressive histone marks H3K9me3 and H3K27me3 are not reduced at ALR in naïve compared to primed hESCs (Supplemental Fig. 2E). In fact, H3K27me3 is higher in naïve than primed hESCs, in agreement with previous reports^38–40^. Of note, the very high expression and chromatin enrichment of ALR RNA in naïve hESCs (Fig. 3D) is specific to ALR, as it is not observed for other types of repeats, including: other human satellite repeats families (BSR/Beta, HSATI, HSAT4, HSAT5), LINE1 families (L1HS, L1PA2, L1PA3, L1PA4) or representative cleavage-stage expressed transposable elements (MLT2A1, MLT2A2, LTR7B, LTR12) (Supplemental Fig. 3A).

We used qRT-PCR to independently validate the strong upregulation of ALR RNA in naïve hESCs, relative to primed hESCs, fibroblasts and HEK293T cells (Fig. 3E). The sharp difference in expression of ALR between naïve and primed hESCs led us to explore a transition model, where naïve hESCs are converted to the primed state by progressively changing the medium (Fig. 3F, top). We found that expression levels of ALR are strongly downregulated as early as 24 hours in full primed cell medium and continue to decrease at 48 hours (Fig. 3F, bottom left). This conversion method is validated by the downregulation of naïve state marker KLF2^35,41^ and upregulation of the primed state marker COL1A1^42^ (Fig. 3F, bottom right). Thus, ALR RNA expression is a novel and specific marker of the naïve state of hESCs.

### ALR RNA localizes to the perinucleolar compartment in naïve hESCs

We next sought to investigate the sub-nuclear localization of ALR RNA in naïve hESCs. Single-molecule RNA Fluorescent In Situ Hybridization (smFISH) revealed a punctate pattern of accumulation of ALR RNA in the nuclei of the naïve hESCs (Supplemental Fig. 4A). The signal is specific to RNA, as it is abolished by RNase A treatment (Supplemental Fig. 4A). This pattern, as expected, is absent in primed hESCs (Supplemental Fig. 4B). We have recently shown that centromeric DNA sequences like ALR are enriched in nucleolar-associated domains in naïve and primed hESCs^24^, in agreement with data in HeLa cells^43^. Given that caRNAs are often retained in cis in close proximity to the DNA elements from which they are transcribed, we explored the localization of ALR RNA foci relative to the nucleolus. RNA FISH-IF revealed that ALR foci are mostly located at the periphery of the nucleolus in naïve hESCs (Fig. 4A). These findings were validated in two other media^24,44^ for naïve hESC culture (Supplemental Fig. 4D). The close association of ALR foci with the periphery of the nucleolus was confirmed using cross-linked RNA immunoprecipitation (CLIP)-qRT-PCR with an antibody against Nucleolin (NCL) (Fig. 4B). NCL is an integral component of the granular compartment, located at the periphery of the nucleolus.

**FIGURE 4.**
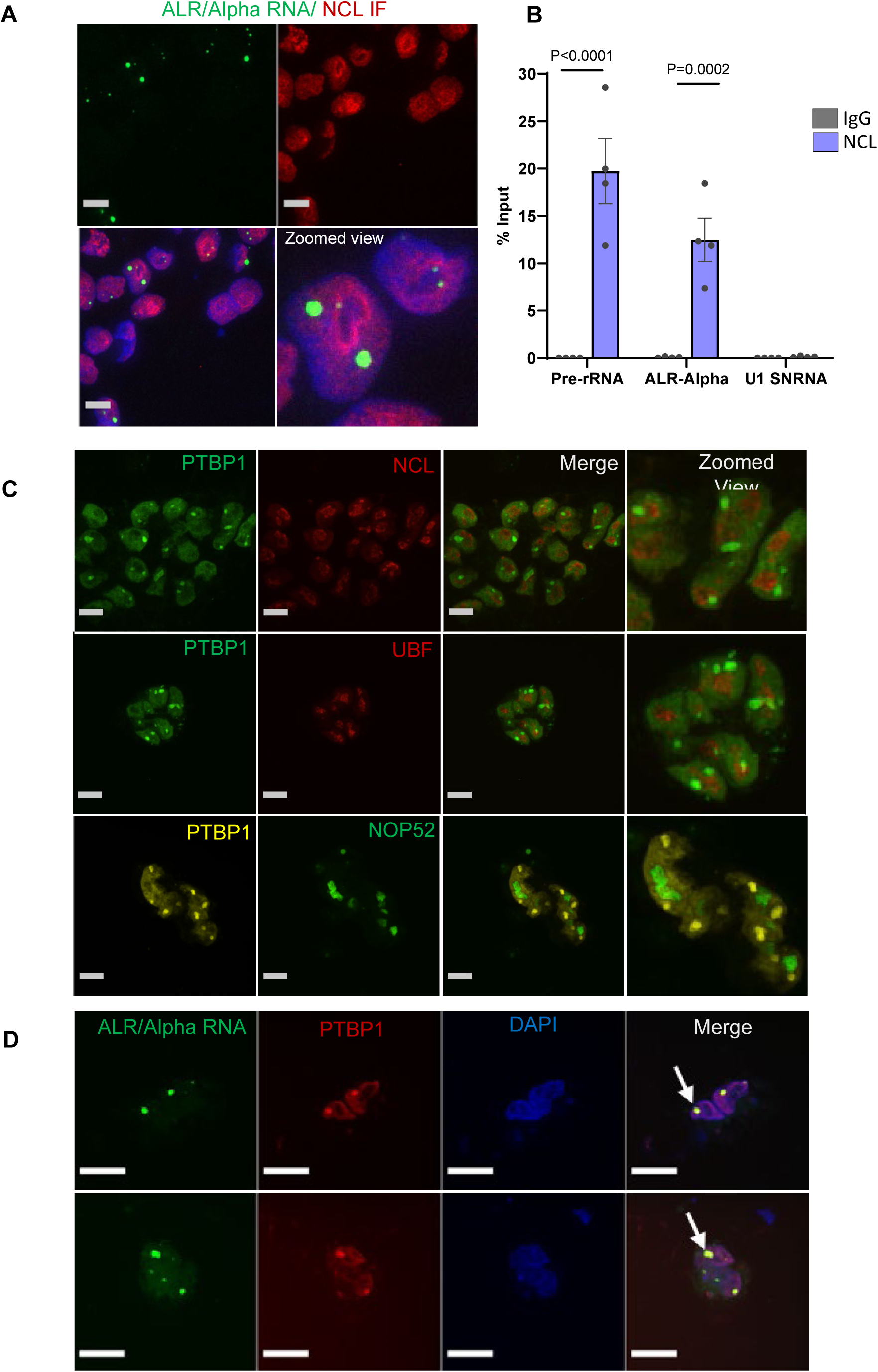
ALR RNA localises to the perinucleolar compartment in naïve hESCs. A) Representative image showing ALR RNA FISH (green) and NCL IF (red) in naïve hESCs. Scale bar, 10 µm. Zoomed view shows proximity of ALR RNA signal to the nucleolus. B) RIP-qPCR in naïve hESC with NCL or control IgG antibodies, showing association of NCL with ALR RNA. Data are n=4 independent experiments. P-values were calculated by two-way ANOVA with Bonferroni’s multiple comparisons test. C) PTBP1 immunofluorescence with antibodies staining nucleolus (NCL, UBF, NOP52) shows perinuclear localization of the PTBP1 in naïve hESCs. Scale bar, 10 µm. D) Representative images showing co-localization of ALR RNA and PTBP1. Cells were stained by IF for PTBP1 combined with ALR RNA FISH. PTBP1 foci are indicated with white arrows. Scale bar, 10 µm.

The nucleus is sub-divided into several compartments that perform distinct functions^45^. We considered whether the ALR RNA foci correspond to a particular compartment. While our data indicate an association between ALR RNA and the nucleolus, the RNA FISH-IF results indicates that it is not localized at the nucleolus. We therefore explored the perinucleolar compartment (PNC)^46,47^. The PNC is defined by the perinucleolar localization of PTBP1 foci. The function of the PNC remains elusive, although it has been implicated in cellular transformation and tumorigenesis. This is in part because the PNC was found to be present only in cancer cells and absent in all normal cells tested, including primed hESCs^31,46,48^. However, we found that distinct PTBP1 foci are detected at the periphery of the nucleolus in naïve hESCs (Fig. 4C), but not in primed hESCs (Supplemental Fig. 4C). Interestingly, RNA FISH-IF revealed a strict co-localization between ALR RNA and PTBP1 protein in naïve hESCs (Fig. 4D). Moreover, ALR RNA and PTBP1 mRNA display similar patterns of expression during early human development, with the highest levels between the 8-cell and blastocyst stages (Supplemental Fig. 4F, G). Thus, the PNC is surprisingly detected in naïve hESCs and is the site of accumulation of ALR RNA.

### Knockdown of ALR RNA induces loss of the PNC and upregulation of rRNA in naïve hESCs

The specific enrichment of ALR RNA in the naïve state of hESCs prompted us to explore its potential function. We designed anti-sense locked nucleic acid gapmers to knockdown (KD) ALR RNA (Fig. 5A, Supplemental Fig. 5A, TableS1). Knockdown of ALR RNA using two independent gapmers was validated by qRT-PCR and smFISH (Supplemental Fig. 5B-D). ALR RNA KD does not appear to have any overt effect on naïve hESC propagation (Supplemental Fig.5E).

**FIGURE 5.**
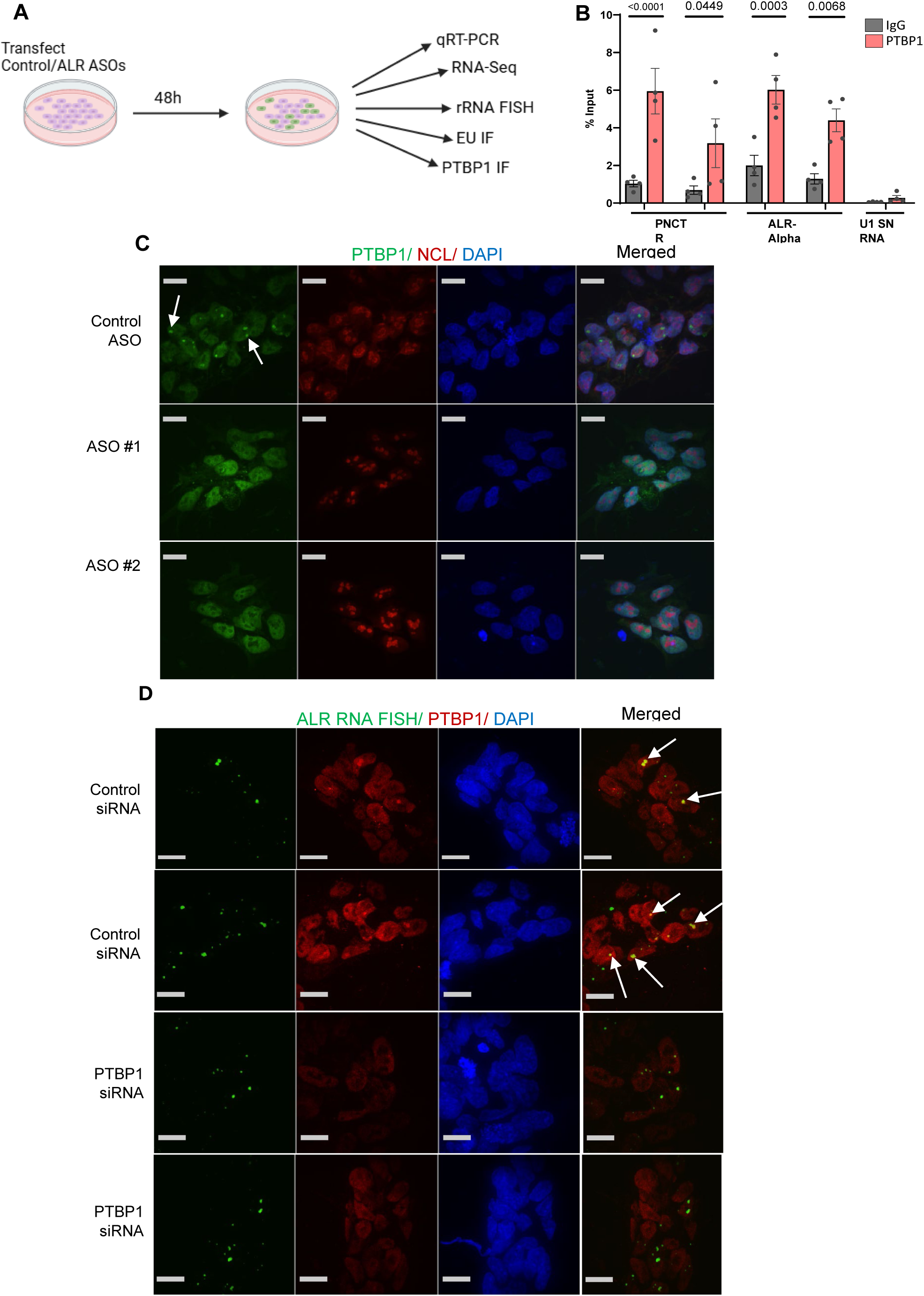
Knockdown of ALR RNA induces loss of the PNC in naïve hESCs. A) Diagram of ALR knockdown and follow-up experiments. B) Cross-linked RNA immunoprecipitation (CLIP) in wild-type naïve hESCs with PTBP1 or control IgG antibodies, showing association of PTBP1 to positive control PNCTR lncRNA and ALR RNAs but nonsignificant association with U1 snRNA, a highly abundant nuclear RNA. Data are n=4 independent experiments. P-values were calculated by two-way ANOVA with Bonferroni’s multiple comparisons test. C) Representative images showing PTBP1 and NCL immunofluorescence in naïve hESC transfected with control ASO or ALR KD ASOs. Cells were stained by IF for PTBP1 and NCL 48h after transfection. PTBP1 perinucleolar foci are lost upon ALR KD (ASO#1 and ASO#2) compared to control ASO. Scale bar, 10 µm. D) Representative images showing ALR-Alpha RNA FISH and PTBP1 immunofluorescence in naïve hESC transfected with control siRNA PTBP1 siRNA. Cells were stained sequentially by FISH for ALR-Alpha RNA and then IF for PTBP1 after treatment with siRNAs for 48h. ALR-Alpha signal is unaffected by PTBP1 KD (PTBP1 siRNA) compared to control siRNA. Scale bar, 10 um.

We first assessed a potential relationship between ALR RNA and the PNC. Using CLIP-qPCR, we found that ALR RNA associates with PTBP1 in hESCs (Fig. 5B), suggestive of a retention of the ALR RNA at the PNC. Interestingly, PTBP1 foci entirely disappear upon ALR RNA KD in naïve hESCs (Fig. 5C). Although ALR RNA KD disrupts the specific localization of PTBP1 in the PNC, the signal for the PTBP1 across the nucleoplasm is unaffected. Given that PTBP1 is a defining factor of the PNC^46,47^, we next explored a potential role of PTBP1 in ALR RNA localization at the PNC. Interestingly, the localization of ALR RNA is unaffected upon KD of PTBP1 (Fig. 5D, Supplemental Fig. 5F). These findings indicate that ALR RNA is required for the compartmentalization of PTBP1 at the PNC in naïve hESCs. In addition, these results suggest that ALR RNA sits upstream of PTBP1 in the organization of the PNC in hESCs.

Given the close association of ALR RNA and the PNC with the nucleolus, where rRNA is transcribed by RNA Pol I, we assessed the levels of rRNA expression. We found that ALR RNA KD leads to a consistent upregulation of rRNA transcription, as assessed by smFISH using probes for 5’ External Transcribed Sequence (ETS) rRNA (Fig. 6A, B). To further explore the role of ALR in rRNA expression control, we directly quantified nascent transcription at the nucleolus. 5-ethynyluridine (EU) incorporation at the nucleolus, a measure of RNA Pol I activity, is remarkably increased upon ALR RNA knockdown, compared to controls (Fig. 6C, D). Both gapmers designed to knockdown ALR RNA consistently lead to significantly increased levels of 5’ETS rRNA and EU, albeit to varying degrees. Overall, these results indicate that ALR RNA, directly or indirectly via the PNC (see Discussion), contributes to repressing rRNA expression in naïve hESCs.

**FIGURE 6.**
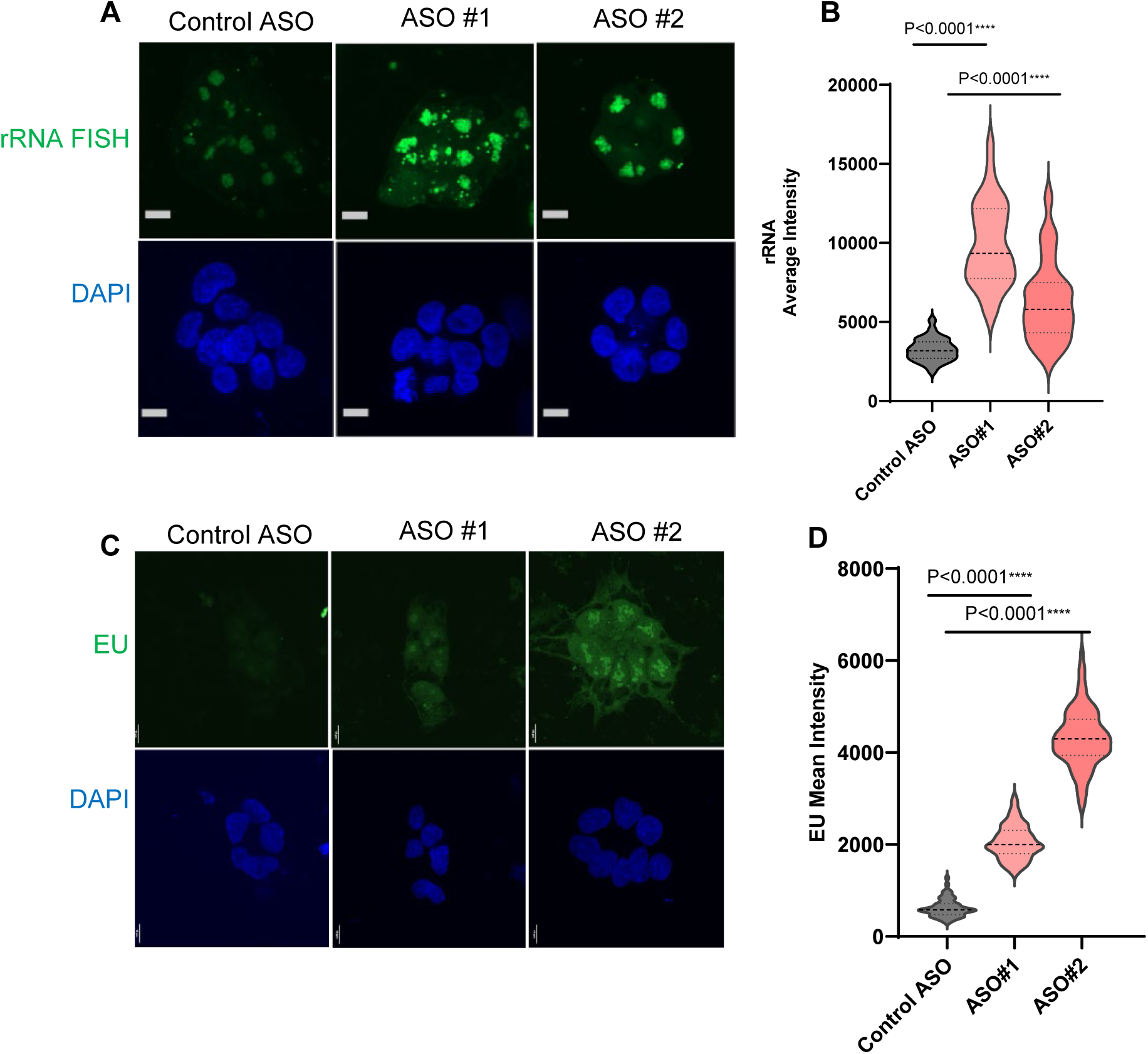
Knockdown of ALR RNA induces upregulation of rRNA in naïve hESCs. A) Representative images of 5’ETS ribosomal RNA FISH 48h after ASO transfection. ALR KD induces reproducible increases in rRNA. Scale bar, 10 µm. B) Quantification of the 5’ETS ribosomal RNA FISH signal. Data are the mean values ± SEM of average FISH signal intensity per experimental batch, n=3 independent experiments. P values calculated by unpaired Student’s t-tests with Welch’s correction. C) Representative EU incorporation-immunofluorescence images 48h after ASO transfection. ALR KD induces reproducible increases in EU incorporation. Scale bar, 10 µm. D) Quantitation of nucleolar EU mean intensity upon ALR KD. Data are the mean values ± SEM of average signal EU signal per experimental batch, n=3 independent experiments. P values calculated by unpaired Student’s t-tests with Welch’s correction.

This repressive role of ALR RNA on rRNA expression in naïve hESCs prompted us to explore their relative expression levels during early human development. In agreement with our findings, ALR RNA and rRNA expression display an overall pattern of anti-correlation in vivo: the upregulation of ALR at the 8-cell and morula stages corresponds to the lowest levels of detection of rRNA (Supplemental Fig. 4F, H). Moreover, chemical inhibition of RNA Pol I in naïve hESCs leads to a strong upregulation of ALR RNA levels (Supplemental Fig. 4I), similar to what has been described in HeLa cells^49^. Taken together, these results indicate that ALR RNA and rRNA have a mutually antagonist relationship in naïve hESCs and early human development.

### ALR RNA KD induces hypertranscription in naïve hESCs

The sharp rise in EU incorporation upon ALR RNA KD described above is not limited to the nucleolus but is rather observed throughout the nucleoplasm (Fig.6C, D). A global rise in EU incorporation and nascent transcription is one of the signs of hypertranscription, the upregulation of the majority of the transcriptome that has been documented in stem and progenitor cells^50–52^. However, EU data does not provide information regarding how many, and which genes may be upregulated. While hypertranscription is masked using standard RNA-seq methods, it can be detected via cell-number normalized RNA-seq (CNN RNA-seq). This involves using the same number of cells in each sample and adding exogenous synthetic RNAs for subsequent normalization^50,53^. We therefore performed CNN RNA-seq in control vs ALR RNA KD naïve hESCs. The results reveal a remarkable upregulation of gene expression across the transcriptome (Fig. 7A, B). Analyses of functional annotations confirmed that ALR RNA KD induces an activation of cellular pathways typically associated with hypertranscription^50,54^, including Myc and E2F targets (Fig. 7C,E) and ribosome biogenesis (Fig. 7D,E). These data indicate that ALR KD in naïve hESCs induces a broad upregulation both of rRNA and mRNAs.

**FIGURE 7.**
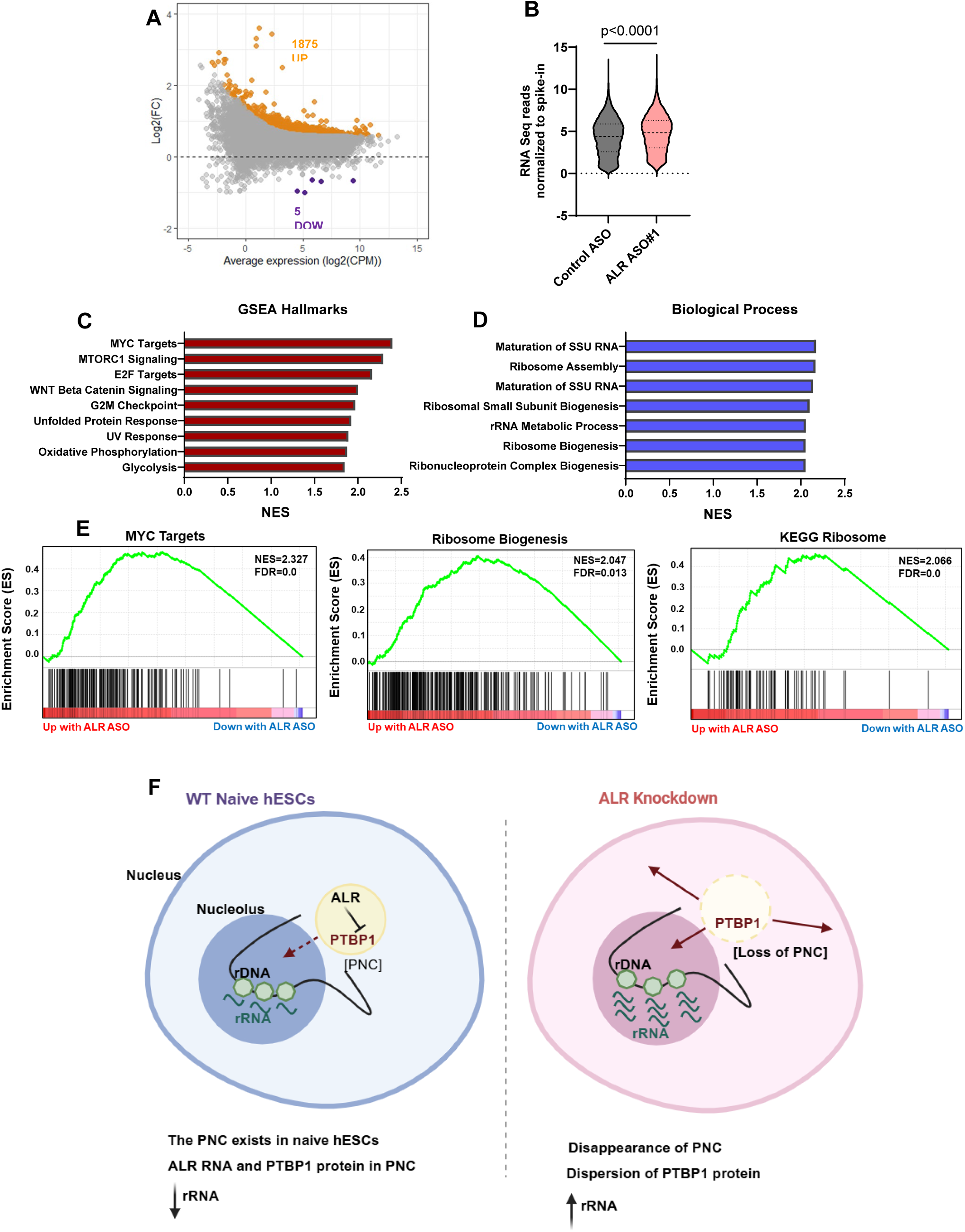
ALR RNA KD induces hypertranscription in naïve hESCs. A) MA plot of CNN RNA-seq data for ALR KD, revealing a global transcriptional upregulation. B) Total reads from RNA-Seq mapping to the human genome normalized to the spike-in reads. P values calculated by unpaired Student’s t-tests with Welch’s correction. C) Gene Set Enrichment Analysis (GSEA) for Cancer Hallmarks shows enrichment of pathways associated with hypertranscription upon ALR KD (MTC targets, MTORC1 signaling, E2F Targets, and others). D) GSEA for biological processes reveals enrichment of hallmarks for ribosome biogenesis pathways upon ALR KD. E) Examples of pathways enriched following ALR KD. F) Model for the role of ALR RNA in naïve human ESCs. We carried out a comprehensive analysis of the cytoplasmic and chromatin associated transcriptomes in human cells, with a focus on ESCs. We found that ALR RNA is highly expressed in naïve but not primed human ESCs and is strongly associated with chromatin. ALR RNA is co-localized with PTBP1 at the perinucleolar compartment (PNC), which was previously thought to only be present in cancer cells. Our results indicate that ALR RNA is required for PNC maintenance and for attenuating global transcriptional output, including of ribosomal RNA, in naïve human ESCs. We propose that ALR RNA sequesters proteins such as PTBP1 in the PNC, preventing from carrying out their roles in transcription and RNA processing, including of rRNA. The results have important implications for our understanding of the role of repeat elements during human development and raise new and testable hypothesis for the potential function of the PNC in cancer progression.

## Discussion

Satellite repeats were first identified in 1960s as distinct bands of DNA that differ from the rest of the genome on a cesium-chloride density gradient^55,56^. In humans, satellite repeats make up about 5-10% of the genome^57–59^. Despite being among the first human DNA sequences to be biochemically isolated and characterized, satellite DNA remains poorly understood. In this study, we carried out a comprehensive characterization of the chromatin-bound transcriptome of different states of hESC pluripotency and identified ALR RNA as a caRNA highly expressed specifically in naïve hESCs. We found that ALR RNA localizes to the PNC, a nuclear compartment marked by PTBP1 previously thought to exist only in cancer cells. Functional data revealed that ALR RNA regulates maintenance of the PNC and global transcriptional state of naïve hESCs. Taken together, our results indicate that ALR RNA functions to maintain nuclear compartmentalization and tune rRNA output during early human development (Fig. 7F), opening new avenues for further exploration.

Satellite repeats generally form condensed constitutive heterochromatin and are kept transcriptionally silent, in part via DNA hypermethylation^60,61^. However, different types of satellite repeats are expressed during early development, cellular stress and in cancer cells^29,50,51,52^. In mouse, major satellite repeats are expressed during early development, with a peak in expression during the 2-cell stage (equivalent to the 8-cell stage in human) and knockdown of their transcripts results in developmental arrest^65–67^. In mouse ESCs, major satellite repeat RNA regulates heterochromatin dynamics and genome stability.^68^ These studies, and the findings reported here, suggest that a more immature state of heterochromatin during development may be a permissive environment for the expression of ALR. Of note, mouse and human naïve ESCs and pre-implantation embryos have overall low levels of DNA methylation, which increases sharply after implantation^69–73^. It will be of interest to determine the role of DNA methylation and other repressive chromatin layers in the regulation of ALR expression in naïve hESCs versus more advanced states of pluripotency and differentiated derivatives. Such studies may shed light on the abnormal de-repression of ALR observed in several types of cancer^62–64,74^.

Our results reveal a surprising role for ALR RNA in attenuating transcriptional output, notably of rRNA, in naïve hESCs. The levels of ALR RNA and rRNA are anti-correlated in hESCs and during early human development, and KD of ALR RNA leads to a sharp increase in nascent transcription of rRNA. This relationship appears to be one of mutual antagonism, as inhibition of RNA Pol I transcription in naïve hESCs (this study) and cancer cell lines^49^ de-represses ALR expression. The mechanism by which, directly or indirectly, ALR RNA regulates rRNA transcription awaits further investigation, but it is likely related to the peculiar localization of ALR RNA. Indeed, an additional contribution of our study is the observation that ALR RNA localizes to the PNC and is required for its maintenance in naïve hESCs. The PNC is a distinct nuclear compartment defined by the focal localization of PTBP1 protein at the periphery of the nucleolus^46,47^. The PNC is detected in various types of cancer cells, and its presence is correlated with disease progression, poorer prognosis and metastases. Moreover, chemically targeting the PNC has shown promise in suppressing metastatic development in several cancer models^75^. Interestingly, the PNC has been claimed to be exclusively detected in cancer cells and to absent in normal cells, including hESCs^31^. Our findings lead us to revise this claim: we confirm that PNCs are absent in primed hESCs, as previously reported, but they are robustly detected in naïve hESCs. Given these findings, we speculate that, in a cancer context, the PNC may endow cells with a more naïve-like state that may favour progression and metastasis.

The function of the PNC remains unclear, although it has been proposed to at least in part act to sequester regulators of RNA biology, including PTBP1, that have roles in splicing, mRNA stability or translation^47,48,76,77^. In mouse ESCs, PTBP1 regulates splicing of hundreds of genes, such as DNMT3B^78^, which then can have a broad impact on gene expression, including as a repressor of rDNA^79^. PTBP1 may also act as transcription factor and bind directly to promoter DNA^80–82^. PTBP1 is significantly overexpressed in various cancers and has been shown to act as an oncogene^32,83–85^. The expression of *PTBP1* during early human development is similar to ALR RNA, and knockout of the mouse *PTBP1* gene leads to early embryonic lethality, prior to gastrulation^46,80–82,86^. Interestingly, PTBP1 has recently been shown to promote rRNA processing and ribosome biogenesis in hematopoietic stem and progenitor cells^87^. Taken together, our results suggest a model (Fig. 7F) in which ALR RNA is a structural component of the PNC that is required for its maintenance in naïve hESCs. We propose that, by sequestering proteins such as PTBP1 at the PNC, ALR RNA contributes to tuning down transcriptional output, including of rRNA, until later stages of differentiation, when ALR transcription is sharply downregulated, and the PNC disappears. Consistent with this model, the KD of ALR leads to dispersion of PTBP1 from the PNC into the nucleoplasm and to a global upregulation of rRNA, which may be in part mediated by PTBP1^87^. At present it is unclear whether the hypertranscription of mRNAs observed upon ALR RNA KD is a secondary effect of the upregulation of rRNA or is directly due to the release of PTBP1 or other regulatory proteins upon dissolution of PNCs. It should prove insightful to identify the protein and RNA content of the PNC in naïve hESCs, and study their function in PNC integrity, transcriptional output and developmental progression. Overall, our findings that PNCs exist in naïve hESCs and depend on ALR RNA provide a new platform to dissect the still elusive cellular and developmental function of this nuclear compartment, shedding light on its potential co-option during cancer progression.

## METHODS

### Ethics statement

All experiments using human ESCs followed the 2021 Guidelines for Stem Cell Research and Clinical Translation released by the International Society for Stem Cell Research (ISSCR). H9 hESC lines are registered for research use in Canada. This research was approved by the Stem Cell Oversight Committee of the Canadian Institutes of Health Research.

### Human ESC culture

Primed H9 hESCs obtained from WiCell were cultured in mTeSR Plus Medium (Stem Cell Technologies 05825) on hESC-Qualified Matrigel (Corning Matrigel, VWR CA89050-192) coated plates in the following conditions: 37°C, 5% CO2, 20% O2. Cell passaging was carried out every 4-5 days using ReLeSR (Stem Cell Technologies 05873) according to manufacturer’s instructions.

Naïve hESCs were cultured in RSeT Medium (Stem Cell Technologies 05975) in the following conditions: 37 °C, 5% CO2, and 5% O2 conditions^33^. RSeT naïve hESC were established via conversion from Primed hESC as per the RSeT Medium manual. Experiments in the converted RSeT hESC were performed within ten passages of RSeT culturing. Naïve hESCs were cultured in a validated naïve media formulation, RSeT+DT, which was supplemented with10nM DZNep (Selleck S7120) and 5 nM TSA (Vetec V900931) for 48h in addition to daily medium change (adapted^44^). 4CL-cultured naïve hESCs were generated from primed hESCs and cultured as previously described^44^.

### Culture of other cell lines

Normal human lung fibroblast cell line IMR-90 were cultured in DMEM Medium (Life Technologies 11965092) supplemented with 10% fetal bovine serum (Atlanta Biologicals S12450), and 1% penicillin-streptomycin (Life Technologies 15140122). Immortalized human embryonic kidney cell line HEK293 was cultured in DMEM Medium (Life Technologies 11995065) supplemented with 10% fetal bovine serum (Atlanta Biologicals S12450), and 1% penicillin-streptomycin (Life Technologies 15140122).

### Cellular fractionation

2-3 x 10^6^ cells were harvested, washed with PBS, and collected by centrifugation. Cell pellets were rinsed in 1× PBS and then the plasma membranes were lysed by resuspension in lysis buffer (10 mM HEPES [pH 7.9], 10mM KCl, 1.5mM MgCl2, 24% sucrose, 10% glycerol, 1mM DTT, 0.1% Triton X-100) and centrifuged for 5 min, 4°C, 13,000xg. The supernatant (cytoplasmic fraction) was collected in a separate tube and kept on ice. The nuclei pellet was gently rinsed with lysis buffer, then resuspended in a chromatin extraction buffer (3mM EDTA, 0.2mM EGTA, 1mM DTT) and centrifuged for 5 min, 4°C, 1,700xg. The supernatant (nucleoplasmic fraction) was collected in a separate tube and kept on ice. The chromatin pellet was gently rinsed with cold 1× PBS and then resuspended in TRIzol (Life Technologies 15596026) using a 21G needle. Chloroform was added to the suspension and centrifuged for 15 min, 4°C, 16,000xg. The aqueous phase (chromatin fraction) was collected in a separate tube and kept on ice. Cytoplasmic and Chromatin fractions were then processed for RNA isolation according to the RNeasy Mini Kit (Qiagen 74104) product manual. On-column RNase-free DNaseI digestion (Qiagen 79254) was carried out on all samples to eliminate genomic DNA contamination and RNA was eluted with nuclease free water in a total volume of 30ul. In addition, off column DNase digestion was also performed on all samples (Life Technologies 18068015), to further ensure removal of genomic DNA contamination.

### qRT-PCR

RNA was quantified using Qubit High Sensitivity (HS) kit (Life Technologies Q32852). Synthesis of complementary DNA (cDNA) was performed using the SuperScript IV VILO Master Mix (Life Technologies 11756050), followed by qPCR quantification using PowerUp SYBR Green Master Mix (Life Technologies A25777) on a QuantStudio 5 Real-Time PCR System (Thermo Fisher). Relative expression data represents average fold change ± Standard Error of Mean (SEM) relative to the average of housekeeping genes UBB and GAPDH, of three biological replicates, plotted and statistically analyzed with Prism GraphPad (Version 9).

### RNA-Seq of cytoplasmic and chromatin RNA

All RNA samples were processed with the NEBNext rRNA depletion kit (NEB E6310) to remove all ribosomal RNAs. Recovered total RNA was quantified by Qubit and quality of the RNA was assessed by Fragment Analyzer HS RNA Kit (Agilent DNF-472). Sequencing libraries were prepared with NEBNext Ultra II Directional RNA Library Prep Kit (NEB E7760L), per manufacturer’s instructions. cDNA library quality was assessed by Fragment Analyzer NGS (Agilent DNF-474). Libraries were quantified by Qubit and pooled at equimolar concentration for paired-end 75bp read sequencing on a NextSeq (Illumina) instrument. Sequencing was performed at the Lunenfeld–Tanenbaum Research Institute Sequencing Facility.

Libraries were quality-checked using FastQC and trimmed of Illumina adaptor sequences using Trim Galore! v0.6.6 (Babraham Bioinformatics) and then aligned to the human reference genome (GRCh38 or T2TCHM13v1.1 where indicated) with hisat2 v2.2.1. Gene and repeat counts were obtained using the Rsubread package (v2.6.4) function “featureCounts” with the following options: isPairedEnd = TRUE, requireBothEndsMapped = TRUE, isGTFAnnotationFile = TRUE, useMetaFeature = TRUE. The counts table was then imported into R studio and converted to a DESeq2 object (DESeqDataSetFromMatrix using sample information) for processing, using DESeq2 version 1.36.0. Genes or repeats were then filtered to remove low count (including those with <10 reads across all samples). Data were further analyzed using tidyverse (v.2.0.0) and plotted using ggplot2 (v.3.4.4). Counts normalized by the DESeq2 rlog transformation were used for PCA and heatmaps of gene expression.

### ALR knockdown

Antisense Locked nucleic acids (LNA) gapmers were designed targeting the full length ALR unit (Repbase^89^) using the Qiagen online gapmer design tool. Multiple gapmers were chosen based on the high design score. These gapmers were then tested for efficiency of knockdown, with two independent gapmers selected for further experiments. The position of the 2 gapmers are as shown in Supplementary Fig. 5A. Qiagen Negative Control A Antisense LNA gapmer was used as a negative control (Table S1). Both 5’-FAM labelled and unlabelled versions of gapmers were used in experiments.

Transfection of gapmers were performed in hESC using 4 µl of lipofectamine STEM Transfection reagent (Life Technologies STEM00008), as per manufacturer’s instructions. Cells were trypsinised using TrypLE (Life Technologies 12605010) and plated at a density of 5 x 10^5^ cells per well of a 6-well plate for transfection. Transfections were performed in 200µl of Opti-MEM medium and 25nM of gapmers, in 2mL of RSeT Medium supplemented CEPT cocktail which includes 50nM Chroman 1 (MedChemExp HY-15392), 5 µM Emricasan (Sigma SML2227), Polyamine (Sigma P8483) and 0.7 µM Trans-ISRIB (Sigma 5095840001) for the first 16 hours. Medium was replaced the next morning with RSeT medium. After 48 hours, unless otherwise specified, cells were harvested or fixed for downstream analyses.

### PTBP1 knockdown

siRNAs targeting PTBP1 have been previously described^76^. For PTBP1 KD, ON-Target plus Human PTBP1 (5725) siRNA-SMART pool (L-003528-00-0005) was ordered from Dharmacon. The Dharmacon siGENOME Non-Targeting Control siRNAs (D-001810-10-5) were used as negative control (Table S1). For siRNAs transfection, DharmaFECT 1 (Dharmacon # T-2001-02) was used. The siRNA transfection was performed following the manufacturer’s instructions, similar to above, including the additions of Trans-ISRIB, Emricasan, and Y-27632. 40 nM siRNAs were transfected, per well of a 6-well plate.

### Cell number normalised RNA-Seq of ALR knockdown

Total RNA was extracted from an equal number of cells (∼2 x 10^5^ cells) using RNeasy Micro Kit (Qiagen 74004) with on-column DNase digestion, followed by off-column DNase digestion (Life Technologies 18068015). RNA extracted from equal number of cells was spiked by adding 4ul of 1:10000 diluted External RNAs Control Consortium (ERCC) Spike-in-Mix1 (Life Technologies 4456740) to each sample.

Sequencing libraries were prepared with ribosomal RNA depletion kit (NEB E6310L) and NEBNext Ultra II Directional RNA Library Prep Kit (NEB E7760L), per manufacturer’s instructions. cDNA library quality was assessed by Fragment Analyzer NGS (Agilent DNF-474). Libraries were quantified by Qubit and pooled at equimolar concentration for paired-end 75bp read sequencing on a NextSeq (Illumina) instrument. Sequencing was performed at Lunenfeld–Tanenbaum Research Institute Sequencing Facility.

Libraries were trimmed of Illumina adaptor sequences using Trim Galore! v0.6.6 (Babraham Bioinformatics) and then aligned to the human reference genome (GRCh38) together with ERCC sequences with hisat2 v2.2.1. Gene and repeat counts were obtained using the Rsubread package (v2.6.4) function “featureCounts” with the following options: isPairedEnd = TRUE, requireBothEndsMapped = TRUE, isGTFAnnotationFile = TRUE, useMetaFeature = TRUE. The raw counts were imported into R and normalised to ERCC reads^90^ using EdgeR (v.3.32.1). Data were further analyzed using tidyverse (v.2.0.0) and plotted using ggplot2 (v.3.4.4). GSEA was carried out with the downloaded version of GSEA, with genes pre-ranked by t-values from the differential analysis (KD/control). Gene set collections were downloaded from the Molecular Signature Database (http://www.gsea-msigdb.org/gsea/msigdb/index.jsp).

### Histone ChIP-Seq Analyses

Raw fastq sequencing files of H3K4me3, H3K27ac, H3K9me3 and H3K27me3 ChIP-seq in naïve and primed hESCs were obtained from data generated by the CEEHRC. ChIP-seq data were assessed for quality using FastQC and then aligned to the T2T-CHM13 (v1.1) genome using Bowtie2 (v2.4.5), allowing for multimapping^91^. Reads aligning to the mitochondrial genome, unknown and random chromosomes, and PCR duplicates were removed. SAM files were converted to BAM files and sorted and indexed using Samtools (v.1.17). The resulting filtered and sorted BAM files were converted to Bigwig files using deeptools bamCoverage (v3.3.0) with counts per million (CPM) normalization for visualization in UCSC genome browser on the CHM13v1.1 assembly.

### RNA Fluorescence In Situ Hybridisation

Naïve hESC were grown on sterilized coverslips in 12-well plates. When the cells were ready (either 48h post-transfection, or when 70% confluence was reached), they were washed once in PBS, fixed in 4% paraformaldehyde for 15 min, followed by one PBS wash and two washes with ice cold 70% ethanol. Coverslips were stored in 70% ethanol in parafilm sealed plates at 4 °C for later use.

For RNA-FISH, cells were rehydrated in RNA wash buffer (WB:10% Formamide + 2xSSC in nuclease free water) at room temperature for two 5 min incubations. Coverslips were then transferred to a piece of parafilm in a 15cm dish and incubated face down on RNA-FISH probe solution. Probes for ALR^49^ and 5’ETS ribosomal RNA^24^ were generated by Stellaris (see TableS1), kept in a stock concentration of 12.5 μM and diluted 1:100 in hybridization buffer. To prepare the hybridization buffer, dextran sulfate (Sigma D6001) was first dissolved in DEPC water at 10%, to which 1 mg/ml E.coli tRNA (Sigma 10109541001), 2X SSC, 0.02% BSA, and 2 mM VRC (NEB S1402S) were added. After sterilizing through a 0.02 μM filter, the mixture was aliquoted into 475 μl per tube and frozen at -20 °C. The hybridization buffer was prepared fresh, by adding 75 μl of formamide per aliquot. Samples were incubated in the dark with probe solution, first for 45 min at 37 °C in a hybridization oven, then at 30 °C overnight without sealing the petri dish with parafilm. The next day, the dried coverslips were rehydrated by adding 50-100 μL of WB at the edges. Coverslips were transferred to a 12-well plate with WB for 5 min, followed by a 1-hour incubation in WB with 2.5 μl/ml RNAseOUT (Life Technologies10777019) at 37 °C. Coverslips were mounted onto microscope slides with VECTASHIELD Antifade Mounting Medium with DAPI and then sealed with nail polish. Images were acquired with Leica Quorum spinning disc confocal microscope. Quantification was performed in ImageJ, with total image planes stacked by ‘average intensity’ projection. Nuclei were quantified using the ROI Manager with subtraction of background values.

### Immunofluorescence

Cells were plated as above for FISH, and when ready washed once in PBS and fixed in 4% Paraformaldehyde (PFA) for 8 min. After 2x PBS washes, the coverslips were either stored at 4 °C for later use or used immediately. The fixed cells were permeabilized with 0.25% Triton X-100 in PBS for 2 min on ice. Cells were then washed twice with PBS-T (PBS + 0.1% Tween-20), followed by 30 min block in PBS-T/1% BSA (Sigma A9418) with 10% Donkey serum (Sigma Aldrich D9663) at room temperature. Primary antibody incubation was performed after blocking for 2h in humified chamber. The following primary antibodies were used: NCL 1:100 (Abcam ab22758), PTBP1 1:100 (Invitrogen 32-4800), UBF 1:100 (UBF1 Alexa fluor 594 (Red)) and NOP52 1:100 (Alexa fluor 488 (Green)). After 3 washes in PBS-T, samples were incubated with fluorescence-conjugated secondary antibodies (if warranted) for 1h at room temperature, followed by mounting of coverslips with VECTASHIELD Antifade Mounting Medium with DAPI (Vector Laboratories VECTH1200). Slides were sealed with nail polish after at least 1h incubation in dark. Images were quantified as above for RNA FISH.

### Simultaneous immunofluorescence and RNA-FISH

For experiments with simultaneous RNA FISH and IF (ALR FISH and NCL IF), the RNA-FISH protocol was performed with following modifications. Primary antibodies were added with the probe in the hybridization buffer at 1:100 dilution and incubated in a humified container, in the dark first for 4 hours at 37 °C, then at 30 °C overnight. The following day, coverslips were transferred to a 12-well plate with WB for 5 min, then transferred to a new well containing WB with 2.5 μl/ml RNAseOUT and 1:200 diluted secondary antibodies, followed by a 1-hour incubation at 37 °C. After two 5 min washes in WB (with 0.5 μl/ml RNAseOUT) at room temperature, coverslips were mounted onto microscope slides with VECTASHIELD Antifade Mounting Medium with DAPI and kept at room temperature for at least1 hour before sealing with nail polish.

### Sequential immunofluorescence and RNA-FISH

For experiments with sequential RNA FISH and IF (ALR Fish and PTBP1 IF), the IF protocol was performed with following modifications. After permeabilization, and a wash with PBS, Primary antibodies were added in PBS at 1:100 dilution and incubated in a humified container, in the dark first for 1 hour at room temperature. After three washes in PBS, secondary antibody (1:200 dilution) was added and incubated at room temperature for 1 hour. After three washes in PBS, cells were fixed again in 4% PFA for 10 minutes and then washed with PBS. Then the protocol of RNA FISH was followed on these coverslips.

### EU labelling and immunofluorescence

Naïve hESC cultured in RSeT medium on coverslips were transfected for 48h with indicated gapmer and then were labelled with 1 mM EU for 45 min for nascent transcription and fixed in 4% PFA for 15 min. Following fixation, samples were permeabilized on ice with 0.25% Triton X-100 in PBS for 5 min. EU fluorescence coupling was performed per the manufacturer’s instructions for the Click-iT RNA Alexa Fluor 594 Imaging kit (Life Technologies C10330). After washes in PBS-T, samples were blocked in PBS-T/1% BSA (Sigma A9418) with 10% Donkey serum (Sigma D9663) for 1 h, incubated with a 1:200 dilution of the NCL primary antibody (Abcam ab22758) for 2h in a humidified chamber. Subsequent secondary antibody incubation and mounting with DAPI stain were performed as described above. Images were captured using Leica Quorum spinning disc confocal microscope. Quantification was performed in ImageJ, with totally image planes stacked by ‘average intensity’ projection. Nucleolar signal of EU intensity was quantified using the ROI Manager to manually encircle based on Nucleolin staining. Image quantification data is presented in violin plot of median fluorescence intensity and analyzed with Prism GraphPad (Version 10) using unpaired t-test with Welch’s correction and is representative of three independent experiments.

### RNA polymerase I inhibition

RSeT hESCs were plated at 4 x 10^5^ cells per well of a 6-well plate and switched to RSeT+DT medium the following day. After 24 hours in RSeT+DT conditions, cells were treated with 0.25μM of BMH-21 (Sigma 5099110001), while DMSO was added to control wells. Cells were fixed after 8 hours for immunofluorescence or collected after 24 hours for RNA isolation.

### RNA Immunoprecipitation and qPCR

Nucleolin RNA immunoprecipitation was performed on nuclear extracts from 1×10^7^ RSeT+DT naïve hESC. Prior to extraction, 5 µg Rb anti-NCL (Abcam 22758) or 5 µg Rb anti-IgG (Millipore CS200581) were pre-bound to 50 µl Protein A Dynabeads (Life Technologies 10002D) and incubated for at least 3 hours rotating at 4 °C. Cells were washed once with cold 1x PBS, then in 1x cold PBS cells were placed on ice without lid and irradiated once with 400 mJ/cm^2^. Following irradiation, cells were harvested with 0.25% Trypsin, pelleted, and resuspended in 2mL PBS, 3mL dH2O and 5mL Nuclear Isolation Buffer (1.28M Sucrose, 40mM Tris-HCl pH 7.5, 20mM MgCl_2_, 4% Triton X-100). Cell lysates were kept on ice for 20 minutes, mixing frequently, then spun down 2500xg for 15 minutes in 4°C. The resulting pellet was resuspended in high salt RIP buffer (1M NaCl, 50mM Tris pH7.4, 1mM EDTA, 0.5% Sodium deoxycholate, 1% NP40, protease and RNase inhibitors) with 0.5mL per IP. Lysates were passed through a 25G1 needle 10 times and centrifuged for 10 minutes at 13000rpm at 4°C. Nuclear lysates were blocked for 30min with 20 µl of Protein A Dynabeads at 4 °C, 30 min. Meanwhile, beads pre-incubated with antibody were next collected on a DynaMag (Thermo Fisher Scientific) and re-suspended in High Salt RIP buffer containing 500 ng/mL tRNA (Invitrogen AM7119) and 1 mg/ml RNase-free BSA (Sigma A9418) to block for 30 min at 4°C. IPs were rotated overnight at 4 °C, then the next day were washed twice with 1 ml High Salt Buffer and then twice with 1 ml Proteinase K Buffer (100 mM Tris-HCl pH 7.4, 50 mM NaCl, 10 mM EDTA, DnaseI (Life Technologies 18068015) and RNase inhibitors (Life Technologies 10777019)). Proteinase K Buffer washes were done for 15 min each on 37°C shaker at 800 rpm. IP samples were eluted in 100uL of Proteinase K Buffer with addition of 50µg Proteinase K (Life Technologies 25530049) and 1 hour incubation at 55°C. RIP RNA was extracted from beads using Trizol and standard phenol-chloroform extraction. The aqueous phase containing the RNA was loaded onto RNeasy mini columns (QIAGEN) with 2x volume of 100% ethanol and RNA was purified as described above, including an on-column DNase I treatment. Purified RIP RNA was then used to generate cDNA for qRT-PCR. Primers are listed in Table S1.

### CLIP-qPCR

Plates of RSeT hESCs were washed once with cold PBS, then were placed in cold PBS on ice with the lid open and UV-irradiated once in UV stratalinker 2400 with 400 mJ/cm^2^. Cells were harvested, and nuclear extraction was performed using a standard Abcam protocol (https://www.abcam.com/protocols/nuclear-extraction-protocol-nuclear-fractionation-protocol). Prior to extraction, 5 mg of PTBP1 or control Mouse-IgG antibodies were pre-bound to 50 ml Protein G Dynabeads (Life Technologies # 10003D) and incubated rotating for 3-6 hours at 4°C. Beads were collected on a DynaMag (Thermo Fisher Scientific # 12321D) and re-suspended in High Salt CLIP buffer [1 M NaCl, 50 mM Tris pH 7.4, 1 mM EDTA, 0.5% Sodium deoxycholate, 1% NP40, protease and RNase inhibitors] containing 500 ng/mL tRNA (Invitrogen # AM7119) and 1 mg/ml RNase-free BSA (Sigma Aldrich # A9418) to block for 30 min, then collected and used immediately in RNA immunoprecipitation. Nuclear extracts (10 x 10^6^ cells per IP) were pre-blocked for 30 min with 20 ml of Protein A Dynabeads at 4°C, 30 min, then incubated with antibody-bound blocked beads overnight at 4°C. Beads were washed twice with 1 ml High Salt Buffer and then twice with 1 ml Proteinase K Buffer [100 mM Tris-HCl pH 7.4, 50 mM NaCl, 10 mM EDTA, DnaseI (Life Technologies # 18068015) and RNase inhibitors (Life Technologies #10777019)]. Proteinase K Buffer washes were done for 15 min each on a 37°C shaker at 800 rpm. After washes, beads were resuspended in 100 ml of Proteinase K Buffer and RNA was eluted via addition of 50 mg Proteinase K (Life Technologies # 25530049) and 1 hour incubation at 55°C. RNA was extracted from beads as described above for RIP RNA extraction. Purified CLIP RNA was then used to generate cDNA for qRT-PCR. Primers are listed in Table S1.

## Data and code availability

Sequencing data, including raw reads and processed data (raw counts for genes and repeats), have been deposited on the NCBI Gene Expression Omnibus (GEO) repository (https://ncbi.nlm.nih.gov/geo) and will be accessible upon publication. The data reference series for this study are GSE290362 and GSE290080.

Data sets of H3K4me3, H3K27ac, H3K9me3 and H3K27me3 ChIP-seq in naïve and primed hESCs were generated in partnership with the CEEHRC, a member of the International Human Epigenome Consortium (IHEC), and will be publicly available at the IHEC’s portal (https://epigenomesportal.ca/ihec).The authors declare that all other data supporting the findings of this study are available within the article and its Supplemental Material. Code supporting this study is available at a dedicated GitHub repository (https://github.com/kmittz/ALR).

## Acknowledgments

We thank Brian McStay, Ji-Young Youn and members of the Santos lab for critical reading of the manuscript. We thank all members of Santos lab for their input throughout the project, in particular J. Zhang for guidance on human embryonic stem cell culture, and T. Macrae, E. Collignon and S. McClymont for guidance on bioinformatics analyses. We are grateful to Brian McStay for the NOP52 and UBF antibodies. We are thankful to K. Chan at the LTRI Sequencing Core, A. Bang at the LTRI Flow Cytometry Facility, R. Bielecki, L. Brown at the LTRI Microscopy Facility and M. Kownacka at the LTRI ESC Facility for their core facility support. This work was supported by Canada 150 Research Chair in Developmental Epigenetics and CIHR Project Grant 505756 to M.R.-S.

## Author contributions

M.R.-S. and K.M. conceived the project and designed the experiments. K.M. performed most of the experiments and interpreted the data. L.M. performed CLIP experiments. M.R.-S. supervised the project. K.M. and M.R.-S. wrote the manuscript with input from all authors.

---

**SUPPLEMENTARY FIGURE 1.**
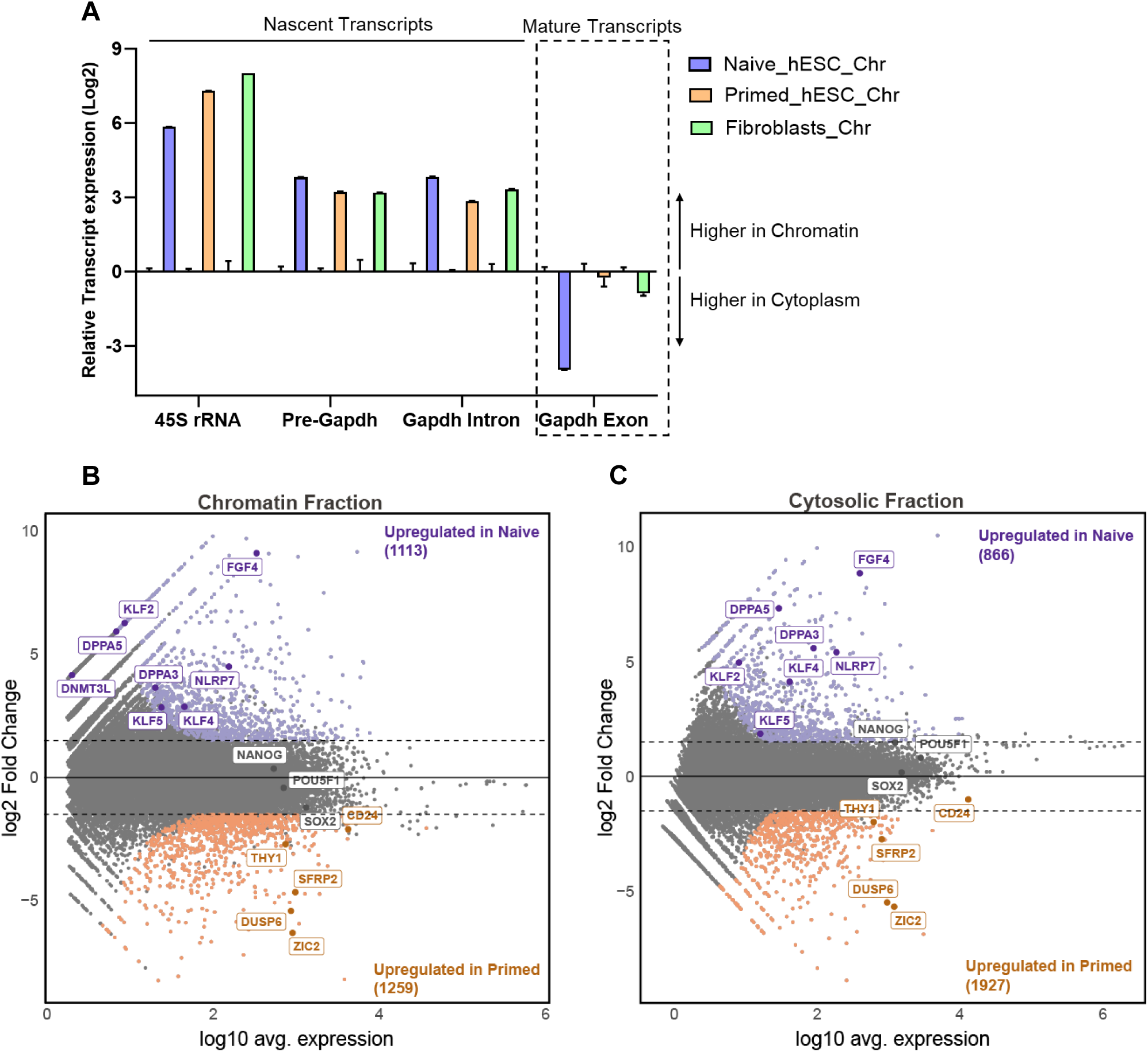
Validation of cellular fractionation, and further comparisons of the transcriptomes of naïve and primed hESCs. A) qRT-PCR to validate the enrichment of nascent transcripts in the chromatin fraction versus the cytoplasmic fraction for naïve hESC, Primed hESC and IMR-90 fibroblasts (n=3). B) MA plot of log2-fold changes in expression between the Naïve chromatin-bound and Primed chromatin-bound conditions. Known markers of naïve hESCs (blue) and primed hESCs (orange) are indicated. C) MA plot of log2-fold changes in expression between Naïve cytoplasmic and Primed cytoplasmic, with similar markers as in B) indicated.

**SUPPLEMENTARY FIGURE 2.**
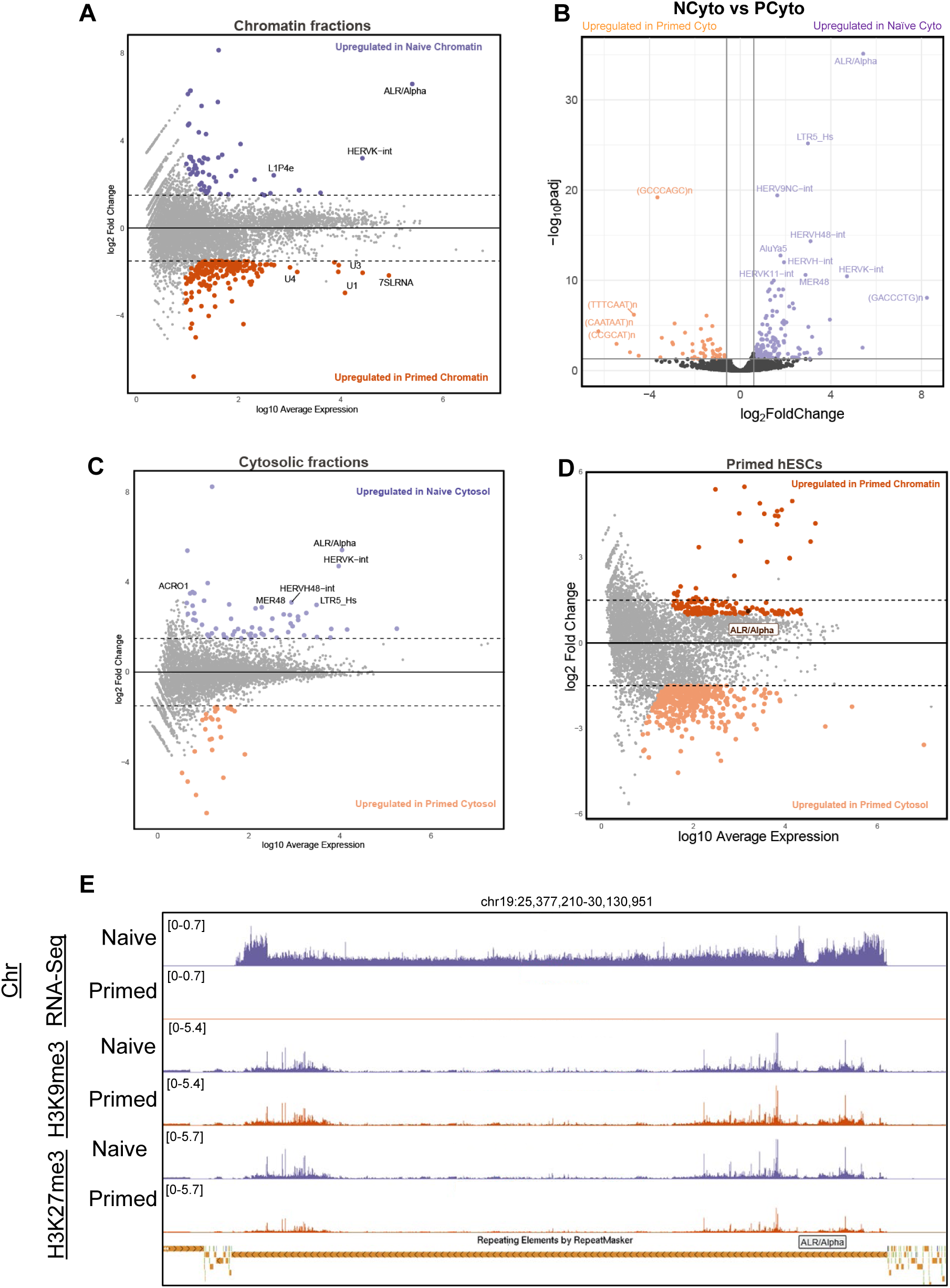
ALR RNA has the highest expression and greatest enrichment at chromatin in naïve hESCs. A) MA plot comparing Naïve chromatin with Primed chromatin fractions. ALR is most highly expressed repeat and is enriched with the highest Log2 fold change in the naïve chromatin fraction. B) Volcano plot comparing Naïve and Primed cytosolic fractions. C) MA Plot comparing Naïve cytosolic with Primed cytosolic fractions. D) MA Plot comparing Primed chromatin with Primed cytosolic fractions. E) Genome browser view of chromosome 19, focusing on the ALR-rich centromere. Chromatin-bound RNA-Seq and ChIP-seq tracks for repressive histone marks (H3K9me3, H3K27me3) in naïve versus primed hESCs are shown (see the Materials and Methods). Tracks from each sequencing experiment were normalized to the same scale between cell types.

**SUPPLEMENTARY FIGURE 3.**
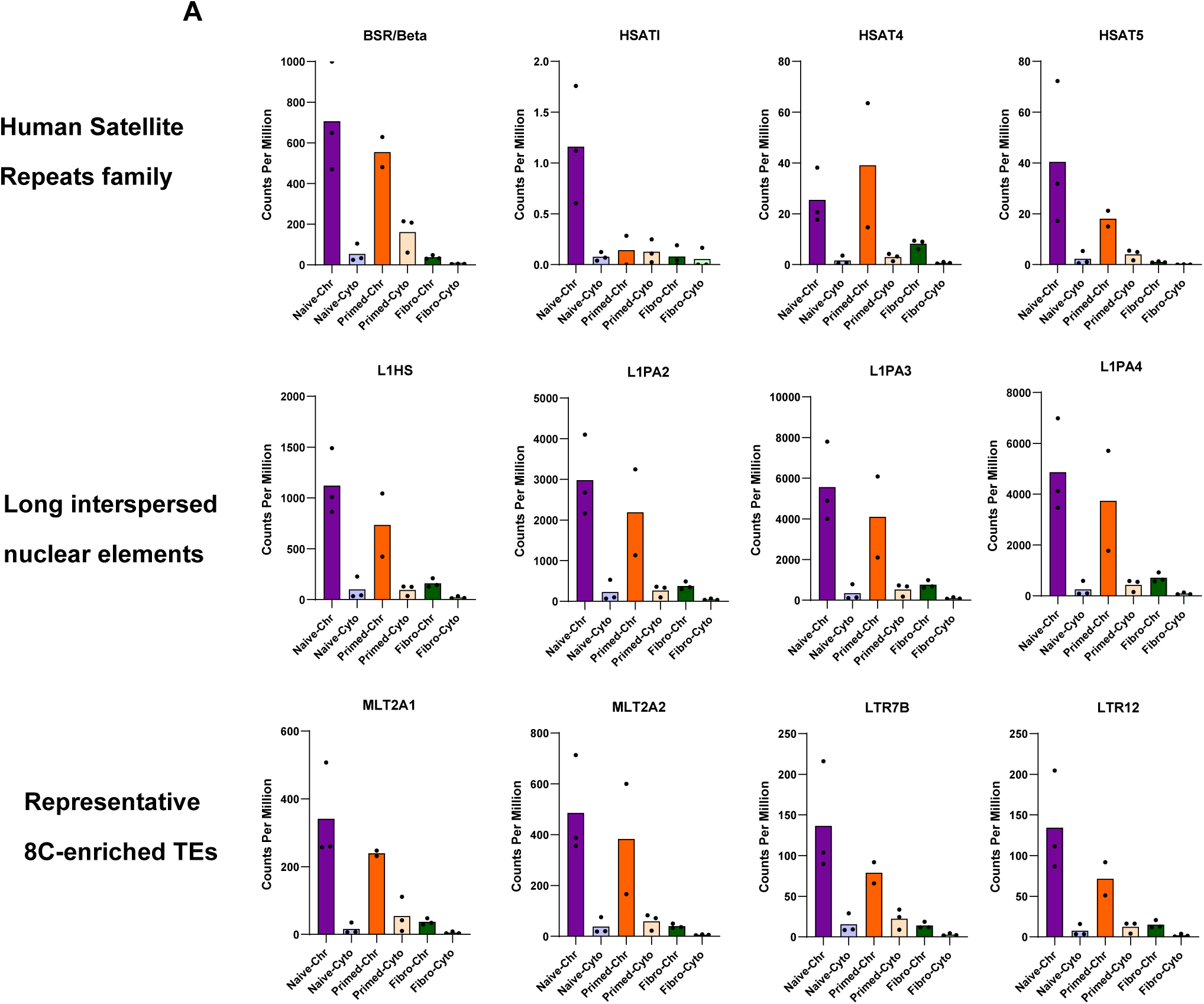
Expression levels of various repeat elements. A) CPM values of indicated repeats from other human satellite repeats, families of long interspersed nuclear element 1 (LINE1), and representative 8-cell enriched TEs do not show the high enrichment in the naïve chromatin fraction observed for ALR (compare to Fig. 3D).

**SUPPLEMENTARY FIGURE 4.**
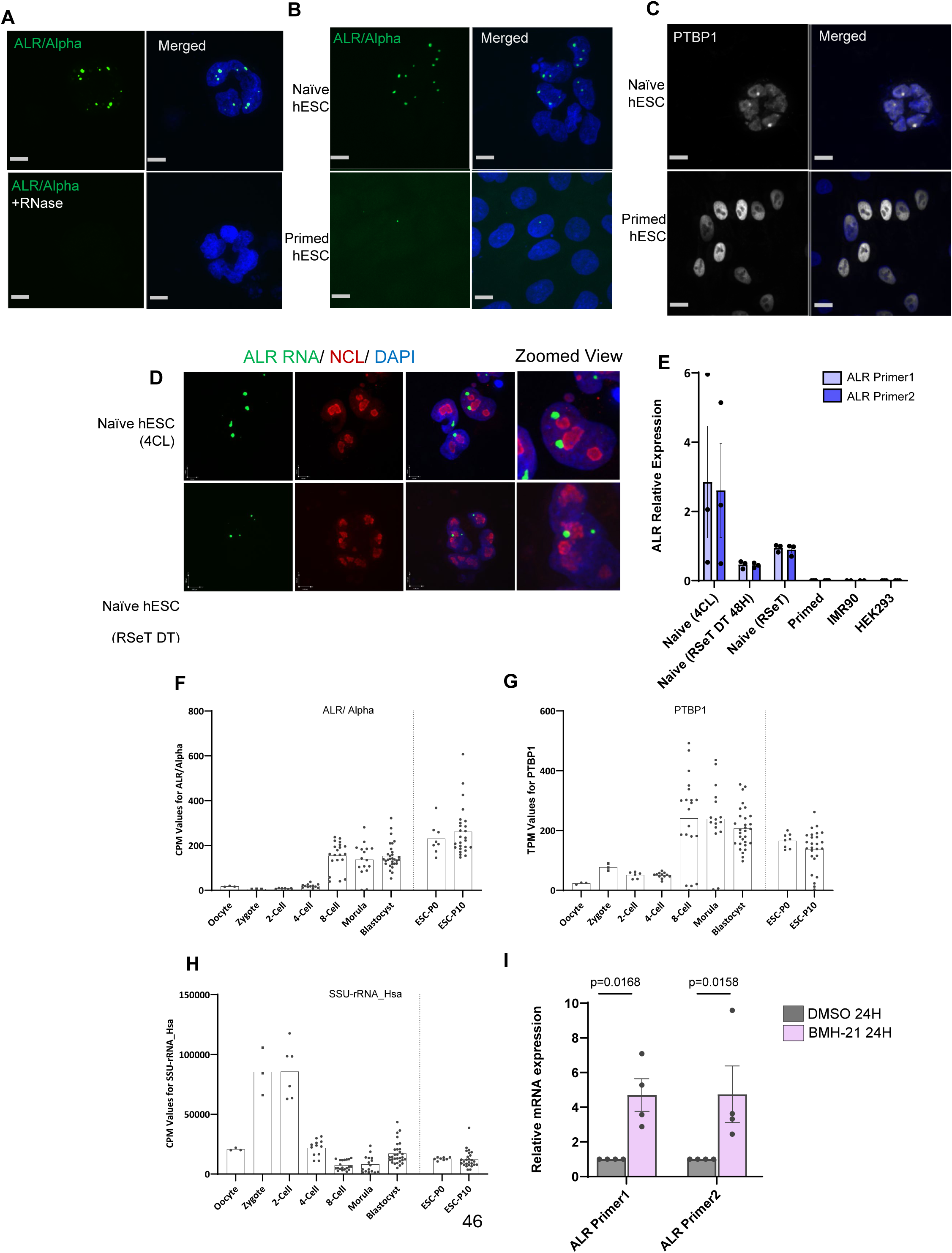
ALR RNA and PTBP1 show punctate localization in naïve hESCs and similar expression profiles during early human embryonic development. A) Representative images showing that smFISH signal for ALR RNA is sensitive to RNase A treatment prior to hybridisation. Scale bar, 10 µm. B) Representative images showing localization of ALR RNA in naïve and primed hESCs. ALR RNA is highly expressed and localized in foci in naïve hESC, but is almost undetectable in primed hESCs. Scale bar, 10 µm. C) Representative images showing localization of PTBP1 in naïve and primed hESCs. PTBP1 foci are present in naïve but not primed hESCs Scale bar, 10 µm. D) ALR RNA FISH (green) and nucleolin immunofluorescence (red) in two independent culture media for naïve hESC (4CL, RSeT+DT). E) Detection of ALR expression in three different naïve hESC culture conditions (RSeT+DT, 4CL and RSeT, see Methods), primed human ESCs, embryonic fibroblasts (IMR-90) and HEK293T cells, by RT-qPCR. Data are average fold change relative to naïve hESC ± SEM, n = 3 biological replicates. F) ALR RNA expression is highest at 8-cell and morula stages of early human embryonic development. Each data point represents the average expression level (CPM) of the ALR in single cells. G) PTBP1 exhibits similar expression patters to ALR during early human embryonic development. Each data point represents the average expression level (TPM) of the PTBP1 in single cells. H) Detection of small subunit rRNA is lowest at 8-cell and morula stages. Each data point represents the average expression level (CPM) of the SSU-rRNA in single cells. In F-H, data were re-analyzed from^30^. I) qRT-PCR for expression of ALR following 24h of PolI inhibition using BMH-21. Data are average fold change ± SEM, n = 4 biological replicates. P-values calculated by two-way ANOVA with Dunnett’s multiple comparisons test.

**SUPPLEMENTARY FIGURE 5.**
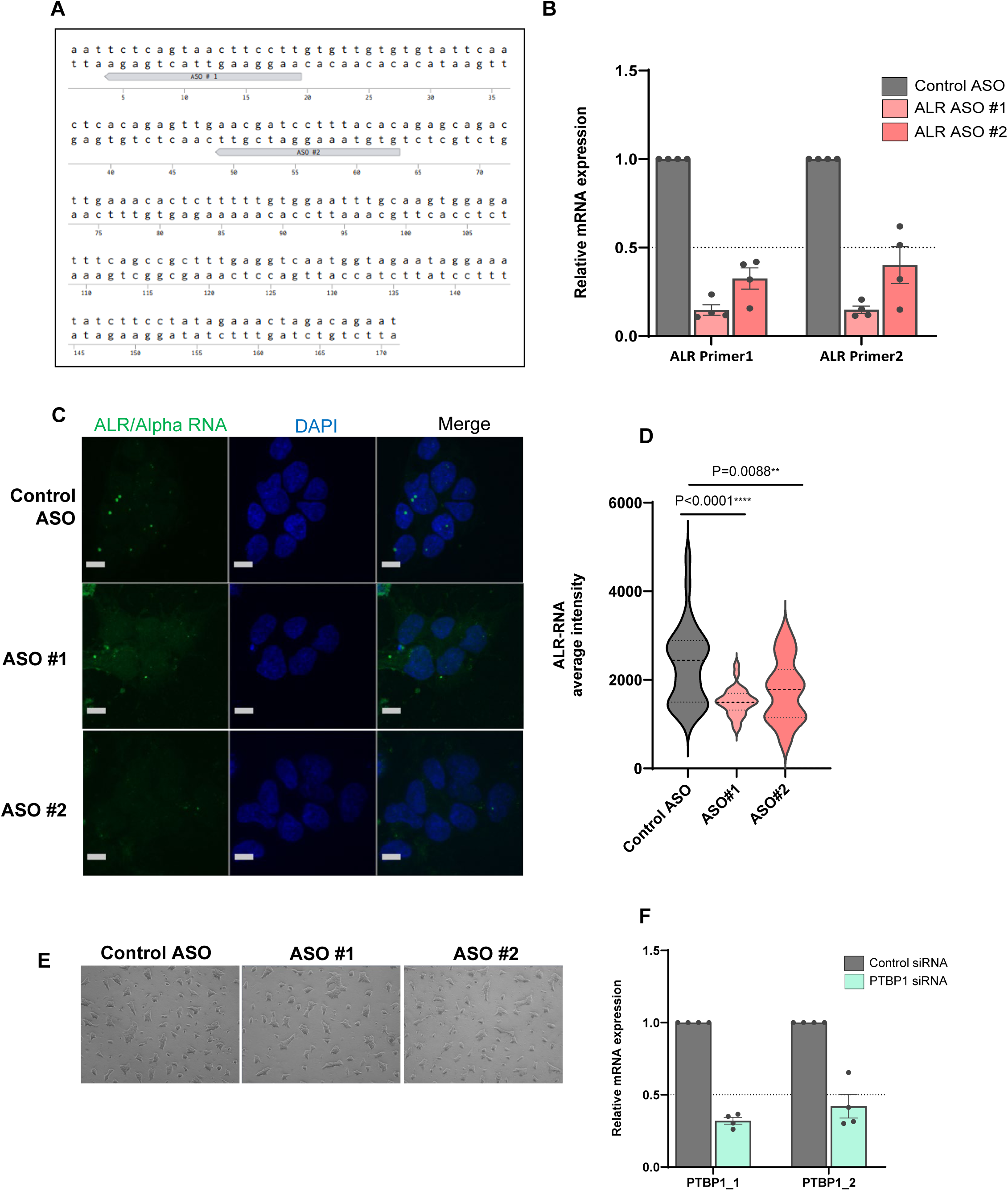
Design and validation of ALR RNA knockdown. A) The ALR consensus sequence was used to design ASOs. ASO#1 and ASO2 are labelled. B) qRT-PCR validation of ALR knockdown following 48h transfection of ASOs. Data are average fold change ± SEM, n = 3 biological replicates. Statistics calculated by two-way ANOVA with Dunnett’s multiple comparisons test. C) Representative images showing ALR in situ hybridisation signal in naïve hESCs after 48h of transfection with control and ASO gapmers. Scale bar, 10 µm. D) Quantification of RNA FISH signal the upon ALR KD. Data are the mean values ± SEM of average signal FISH signal per experimental batch, n=3 independent experiments. P values calculated by unpaired Student’s t-tests with Welch’s correction. E) Representative images of naive hESCs 48h after transfection with ASOs against ALR. F) qRT-PCR validation of PTBP1 knockdown following 48h transfection of siRNAs. Data are average fold change ± SEM, n = 3 biological replicates. Statistics calculated by two-way ANOVA with Dunnett’s multiple comparisons test.

